# Apelin inhibits cyst growth and improves kidney function in mice with polycystic kidney disease

**DOI:** 10.64898/2026.03.26.714294

**Authors:** Duuamene Nyimanu, Anubhav Chakraborty, Stephen C Parnell, Darren P Wallace, Alan SL Yu

## Abstract

**Background:** Autosomal dominant polycystic kidney disease (ADPKD) is a common inherited disorder marked by numerous renal cysts that impair kidney function, with about half of affected individuals progressing to kidney failure by midlife. Patients exhibit reduced circulating apelin, a ligand of the apelin receptor, known to regulate cardiovascular function including hypertension. We tested whether diminished apelin signaling contributes to cystogenesis and if exogenous apelin receptor activation can improve disease outcomes.

**Methods:** Plasma samples from age- and sex-matched healthy controls and ADPKD participants were analyzed for circulating apelin peptides. To assess direct cystic effects, primary ADPKD renal epithelial cells were grown as 3D collagen-embedded cysts and treated with apelin agonists. Male and female *Pkd1^RC/RC^; Pkd2^+/−^* (PKD) mice were treated for 27 days with apelin agonists, vehicle, or the standard of care drug, Mozavaptan. Kidney and heart weight ratios, BUN, renal cAMP, and kidney transcriptional profiles were evaluated.

**Results:** Circulating apelin peptides were significantly reduced in ADPKD patients despite normal kidney function (eGFR, BUN, and creatinine). *In vitro*, both apelin and the small molecule apelin receptor agonist Azelaprag inhibited cyst growth. Apelin and Mozavaptan reduced kidney weight, cystic index, blood urea nitrogen and renal cAMP in PKD mice, whereas Azelaprag did not. Apelin downregulated expression of genes associated with cyst progression, including *Lcn2 (Ngal)*, *Postn,* and *Havcr1 (Kim-1)*. Mozavaptan, but not apelin, induced diuresis and reduced urinary concentration.

**Conclusion:** Apelin receptor activation by exogenous apelin inhibited cAMP synthesis and cyst growth and improved kidney function in an orthologous mouse model of ADPKD. We propose that the apelin receptor may be a potential therapeutic target in ADPKD.

## Introduction

Autosomal dominant polycystic kidney disease (ADPKD) is a common inherited kidney disease affecting at least 12 million people globally^1^. The disease is caused predominantly by mutations in *PKD1* and/or *PKD2*, which encode polycystin-1 and polycystin-2, respectively. Polycystin-1 & 2 are thought to form functional heteromers that regulate intracellular calcium signaling and maintain the differentiated phenotype of renal epithelial cells. Mutations in the PKD genes cause aberrant intracellular calcium regulation, cAMP-driven renal epithelial cell proliferation and transepithelial fluid secretion, resulting in the formation and growth of fluid-filled cysts^2–4^. The progressive growth of cysts leads to nephron loss and massively enlarged kidneys, inflammation and fibrosis, resulting in a decline in renal function. Approximately, one-half of the ADPKD patients progress to end-stage renal disease by the sixth decade of life.

Currently, tolvaptan (Jynarque) is the only approved drug to slow cyst enlargement in ADPKD patients. Tolvaptan is a small molecule antagonist of the arginine vasopressin (AVP) receptor type 2 (V_2_R), a G protein-coupled receptor expressed in the principal cells of the collecting ducts. The binding of tolvaptan to V_2_Rs inhibits AVP-mediated cAMP accumulation, a key driver of cystogenesis through activation of ERK-dependent cell proliferation^4^. Side effects of tolvaptan include polyuria, nocturia, polydipsia and liver toxicity in some patients, limiting its clinical use. One of the major complications of ADPKD is cardiovascular disease, and treatment with tolvaptan does not have any direct cardiovascular benefit^5^. Hence, there remains an urgent need for safe and effective treatment options for these patients.

The apelin system comprises a G protein-coupled receptor (GPCR), the apelin receptor (previously called APJ) and two cognate ligands, apelin and Elabela/Toddler (ELA). It has emerged as an appealing therapeutic target, particularly for cardiovascular diseases (CVD), as it promotes endothelium-dependent vasodilation, decreased blood pressure, and increases cardiac inotropy^6–10^. It is also being investigated as a potential therapeutic agent for metabolic diseases^11^. The apelin receptor primarily signals via the inhibitory Gαi/o protein family, reducing cAMP production. In the kidney, apelin signaling inhibits AVP-mediated water reabsorption by inhibiting cAMP accumulation and aquaporin-2 localization at the apical membrane of collecting ducts^12–14^. In addition, the apelin receptor has been shown to transduce signals via the Gαq protein family and β-arrestin^15,16^.

In animal models of renal ischemia reperfusion injury, mRNA expression levels of apelin receptor ligands were reduced, and exogenous treatment with either apelin or elabela ameliorated injury-induced tubular lesions, renal cell apoptosis, and fibrosis; and improved kidney function (via normalizing urine creatinine and proteinuria)^17–19^. In animal models of cardiovascular disease, activation of the apelin receptor decreased cardiac hypertrophy and fibrosis, inhibited renin-angiotensin system activation, and improved vascular function^10,20^. Recently, phase 1 studies showed AMG986 (also called Azelaprag), a small molecule apelin receptor agonist developed for the treatment of heart failure, to be safe and well-tolerated, even in patients with a loss of renal function^21–24^.

In ADPKD patients, circulating apelin levels were significantly lower than those of healthy subjects and inversely correlated with levels of the copeptin, a surrogate measure of vasopressin release^25–27^. Follow-up evaluation of these patients showed that decreased circulating apelin was independently associated with decrease in estimated glomerular filtration rate (eGFR) and increase in total kidney volume^25^. These data suggest that decreased apelin signaling might elevate intracellular cAMP in collecting ducts, driving cyst growth, and that apelin agonists would be beneficial in ameliorating this disease.

Here, we confirmed that circulating levels of apelin peptides were significantly downregulated in children and young adults with ADPKD and tested the hypothesis that exogenous apelin ameliorates renal cystic disease in an orthologous mouse model of PKD. We also determined if exogenous apelin inhibited cAMP production and signaling, as well as ERK1/2 activation, and *in vitro* cyst growth of primary human ADPKD cells. Our results suggest that the apelin receptor could be a potential novel therapeutic target for treating renal cyst growth in ADPKD.

## Materials and Method

### Human Studies

Plasma samples were collected in EDTA tubes from children and young adults with ADPKD and age- and sex-matched healthy subjects through the multicenter Early PKD Observation Cohort (NCT02936791). Plasma apelin levels were measured using a commercial apelin ELISA kit (cat. No. EK-057-23, Phoenix Pharmaceuticals, Burlingame, USA) according to the manufacturer’s recommendations.

### Animals

*Pkd1^RC/RC^* mice, kindly provided by Peter Harris (Mayo Clinic), and *Pkd2^-/+^* mice, provided by Stefan Somlo (Yale) ^28,29^ were maintained by the PKD Rodent Models and Drug-Testing Core at the University of Kansas Medical Center. *Pkd1^RC/RC^* mice were bred with *Pkd2^+/-^*(both on the C57/6J background) to produce *Pkd1^RC/RC^;Pkd2^+/-^*, an early-onset, rapidly progressive model of polycystic kidney disease (PKD)^30^. Non-cystic littermates (both *Pkd1^+/+^/Pkd2^+/-^*and *Pkd1^+/+^/Pkd2^+/+^*) were used as controls. All the mice used in this study were housed in micro-isolator cages on air-filtered ventilated racks. Animal studies were approved by the Institutional Animal Care and Use Committee at the University of Kansas Medical Center and adhered to the National Institutes of Health Guide for the Care and Use of Laboratory Animals.

### PKD mouse experiments

Mozavaptan (Cat. No.: HY-123593), [Pyr^1^]apelin-13 (Cat No. HY-P1033), and azelaprag (Cat. No. HY-109111) were obtained from MedChemExpress (Monmouth Junction, New Jersey, USA). Male and female animals were randomly assigned to four groups: vehicle, mozavaptan (0.1% in powdered chow^31^, [Pyr^1^]apelin-13 (2 mg/mL body weight), and azelaprag (2 mg/mL body weight), and treated from postnatal day 22-48, with daily intraperitoneal injection of vehicle, [Pyr^1^]apelin-13, and azelaprag. On postnatal day 49 (7 weeks), animals were weighed and blood collected by cardiac puncture under anesthesia. Both kidneys and heart were harvested and weighed. The left kidney was fixed in 4% paraformaldehyde for 24 h followed by storage in 80% ethanol at 4°C until blocking and sectioning for histology. The right kidney was snap frozen and stored at -80°C until recovery for protein and RNA extraction.

### Measurements of cystic index and blood urea nitrogen (BUN)

Kidneys were cut transversely into two halves, processed, and sectioned at the Histology Core at the University of Kansas Medical Center. Sections (5 µm thick) were deparaffinized, rehydrated, and Masson Trichrome staining (Cat. No. NC0017745; Polysciences, Warrington, PA) performed according to the manufacturer’s protocol. Kidney sections were automatically imaged on a Nikon Eclipse High Content Analysis (HCA) system (Tokyo, Japan), comprised of a Ti-E inverted motorized microscope and cystic index quantified as previously described^32^. BUN was quantified using the Quantichrom Urea Assay kit (Bioassay Systems), according to manufacturer’s protocol.

### Measurements of renal fibrosis

Kidney sections were deparaffinized and rehydrated before staining with Picro-Sirius Red (Cat. No. 77455; ScyTek Laboratories, Inc, Logan, UT) according to manufacturer’s protocol. Briefly, Picro-Sirius Red was added to rehydrated sections for 60mins at room temperature, rinsed twice in acetic acid solution (0.5%), and dehydrated in absolute ethanol. Sections were mounted and imaged before quantification of fibrosis area as a percentage of the entire tissue section.

### Measurements of renal cAMP levels

About one-third of a kidney was weighed and homogenized in 10 volumes of 0.1 M HCl (with ∼5 mM IBMX) in a 2 mL glass-Teflon tissue grinder. After centrifugation at 600∈ g for 10 min, the supernatant was assayed using cyclic AMP select Kit according to manufacturer instructions (Cat. No.: 501040, Cayman Chemical, Ann Arbor, MI). Lysates were aliquoted, and protein concentrations were determined using the Pierce 660 nm protein assay kit (ThermoFisher), according to manufacturer instructions. The cAMP concentration was normalized to protein concentration as pmol/mg protein.

### RNA extraction and quantitative reverse transcription-PCR (qPCR)

Whole-tissue RNA was extracted using TRIzol and the PureLink RNA Mini Kit (Invitrogen, Cat #12183018A) according to the manufacturer’s protocol. A high-capacity cDNA Reverse Transcription Kit (Invitrogen, Cat. # 4374966) was used for first-strand cDNA synthesis with 1 μg mRNA. mRNA quantification of the kidney was performed by Taqman assay using QuantStudio™ 5 Real-Time PCR System, 384-well (Applied Biosystems), and primer codes (Table S1). Expression levels were normalized to ezrin, a housekeeping gene as previously described^33^.

### RNA Sequencing

Kidneys were collected from vehicle-treated and apelin-treated male and female mice (n=3 per group) and sent to KUMC Genomics core for total extraction and quality control (for RNA quality/quantity) before library preparation. Pooled libraries were sequenced on Illumina NovaSeqX Plus instrument using standard protocols for paired end sequencing at a depth of 25 million reads per sample.

### Immunoblot analysis

Protein samples were extracted with 0.1M HCl and then neutralized (1:1) in Tris-HCl buffer (50 mM Tris, 5 mM MgCl_2_, 5 mM EDTA, pH 7.4) with protease and phosphatase inhibitors. Samples were mixed with Laemmli buffer containing tris(2-carboxyethyl)phosphine (TCEP) and heated to 95° C for 5 min before loading onto an SDS-polyacrylamide gel. Proteins were then transferred to a nitrocellulose membrane using the TransBlot Turbo semi-dry transfer system (BioRad) and blocked in 5% milk for 30 min at room temperature. Blots were incubated in rabbit anti-phospho-ERK1/2 (1:1000, CST 9101S) or anti-ERK1/2 (1:1000, CST 9102) overnight at 4° C. Horseradish peroxidase-conjugated secondary antibodies were incubated at a 1:10,000 (GE Healthcare Bio-Sciences), and bands were detected by chemiluminescence and quantified.

### Measurement of urine output and osmolality

Wild type mice (24-36 week old) were acclimatized in a metabolic cage for two days with *ad libitum* food and water and returned to their home cage. On day 3, mice were given an intraperitoneal injection of vehicle (0.9% saline) and placed in metabolic cages for four hours before urine was collected, weighed, and centrifuged at 10,000∈ g for 5 min. The following day, mice were split into two groups: one receiving [Pyr^1^]apelin-13 and the other receiving azelaprag and given intraperitoneal injections of two different doses (2 mg/kg BW and 4 mg/kg BW) of [Pyr^1^]apelin-13 and azelaprag on two successive days. Urine was collected as described and stored at -20° C until urine osmolality was determined with a Wescor Vapor Pressure Osmometer (Wescor, Rhode Island, USA).

For mozavaptan, mice (16-44 weeks) were given a powdered diet together with normal chow to acclimatize for five days before they were given only the powdered diet for two days, and on the third day, the food was mixed with water (vehicle) overnight. Mice were placed in metabolic cages, and urine was collected and weighed as before. Then, the mice returned to their home cage with 0.1% mozavaptan solubilized in water and mixed into powdered chow overnight. The following day, mice were placed in metabolic cages for five hours for urine collection as previously done.

### Human primary ADPKD cell culture and microcyst experiment

Primary ADPKD cells were generated by the PKD Biospecimens and Biomaterials Core in the Kansas PKD Center, which is part of the PKD Research Resource Consortium. Cell proliferation assay (see supplementary) and *in vitro* 3D microcyst assay experiments were performed as previously described^30,34^ with some modifications. Briefly, ADPKD cells (9 × 10^5^ cells) were suspended in defined media (1:1 RenaLife Basal Media and Advanced MEM supplemented with 1% PenStrep and ITS, 5 pmol/L triiodothyronine and 50 ng/mL hydrocortisone) and mixed with type 1 collagen (PureCol, Advanced Biomatrix). Immediately after adding collagen, 100 µL cell suspension in collagen was added to each well of a 96 well plate and incubated at 37° C for 45 min to polymerize. Upon polymerization, 150 µL defined media containing 10 µmol/L forskolin and 25 ng/mL EGF was added to each well to initiate cyst growth. Following cyst formation on day 4, the media was changed to 5 µmol/L forskolin and 5ng/mL EGF for 24 h before treatment with [Pyr^1^]apelin-13 (1 nmol/L to 1 µmol/L) or azelaprag (1 nmol/L to 1 µmol/L) in the presence of 5 µmol/L forskolin and 5 ng/mL EGF for 7 days. Treatment reagents were replaced every 2 days for each condition until the end of the experiment. Media were removed and 1% formalin was added to each well to fix cysts in the collagen gels. Plates were imaged using a digital camera attached to an inverted microscope and analyzed with the Image Pro-Premier software (Media Cybernetics). Cyst volume was determined from measurements of the outer diameters (using formulae for volume of a sphere: 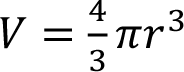 and total cyst volume determined from the sum of individual cysts within each well. Cysts with a volume less than 13 µm were excluded from analysis as previously described^30^. This experiment was also repeated with apelin receptor agonist treatment in the presence of 5 µmol/L forskolin.

## Statistical Analysis

Statistical analyses were performed with GraphPad 10 software, and Student’s 2-tailed t-tests were used to test differences between the two groups after performing a normality test. Where any group was not normally distributed, the corresponding non-parametric statistical test was performed. One- or two-way ANOVA was used to test for differences in multi-group experiments with Dunnett’s multiple comparisons test. *P* values less than 0.05 were considered significant.

## Results

### Expression of apelin signaling in normal and ADPKD patients

Circulating levels of apelin were found to be significantly reduced in adult ADPKD patients compared to healthy controls^25–27^. These previous studies focused on older ADPKD patients with established disease (particularly in Lacquaniti *et al.*^25^), therefore, we evaluated circulating levels of apelin peptides in children and young adults with ADPKD. Plasma samples were obtained from the longitudinal, multicenter Early PKD Observation Cohort (EPOC) study, which included ADPKD patients and age- and sex-matched healthy controls. We found that circulating apelin levels were significantly reduced in ADPKD children and young adults with normal eGFR (Fig. 1A-F; Table 1). Compared to healthy controls, BUN and creatinine concentrations in these patients were not different from those of healthy subjects, and eGFR was slightly higher in the ADPKD patients than healthy subjects (Table 1). Our correlation analysis did not find a significant positive or negative association between circulating apelin levels and creatinine, BUN, or eGFR (Fig. 1C-E), but both age and htTKV showed a positive correlation with apelin levels (Fig. 1B, F). We also analyzed data from ADPKD patients by sex since apelin is an X-linked gene and it has been suggested that females have higher levels than males in certain diseases^35,36^; however, we did not find a sex difference in circulating apelin levels (Fig. S1A). Additionally, there was not any difference in circulating levels of apelin when grouped based on the Mayo Imaging classification (Fig. S1B). To further test the hypothesis that apelin signaling is downregulated in ADPKD kidneys, we analyzed microarray expression data (Song *et al.*^37^) for changes in expression of the genes encoding the apelin receptor (*APLNR*), apelin (*APLN*), and elabela/toddler (*APELA*). *APLN* mRNA (Log2 fold change) was not different between cystic and normal cortical tissues of these patients (Fig. S2C). By contrast, *APLNR* mRNA (Log2 fold change) was lower in medium- and large-sized cysts compared to normal cortex, but it remained unchanged in small cysts or minimally cystic kidneys (Fig. S2A). *APELA* levels appeared to be sensitive to cystogenesis, since there was a significant decrease in mRNA expression in all cyst sizes compared to the normal cortex (Fig. S2B).

**Figure 1.**
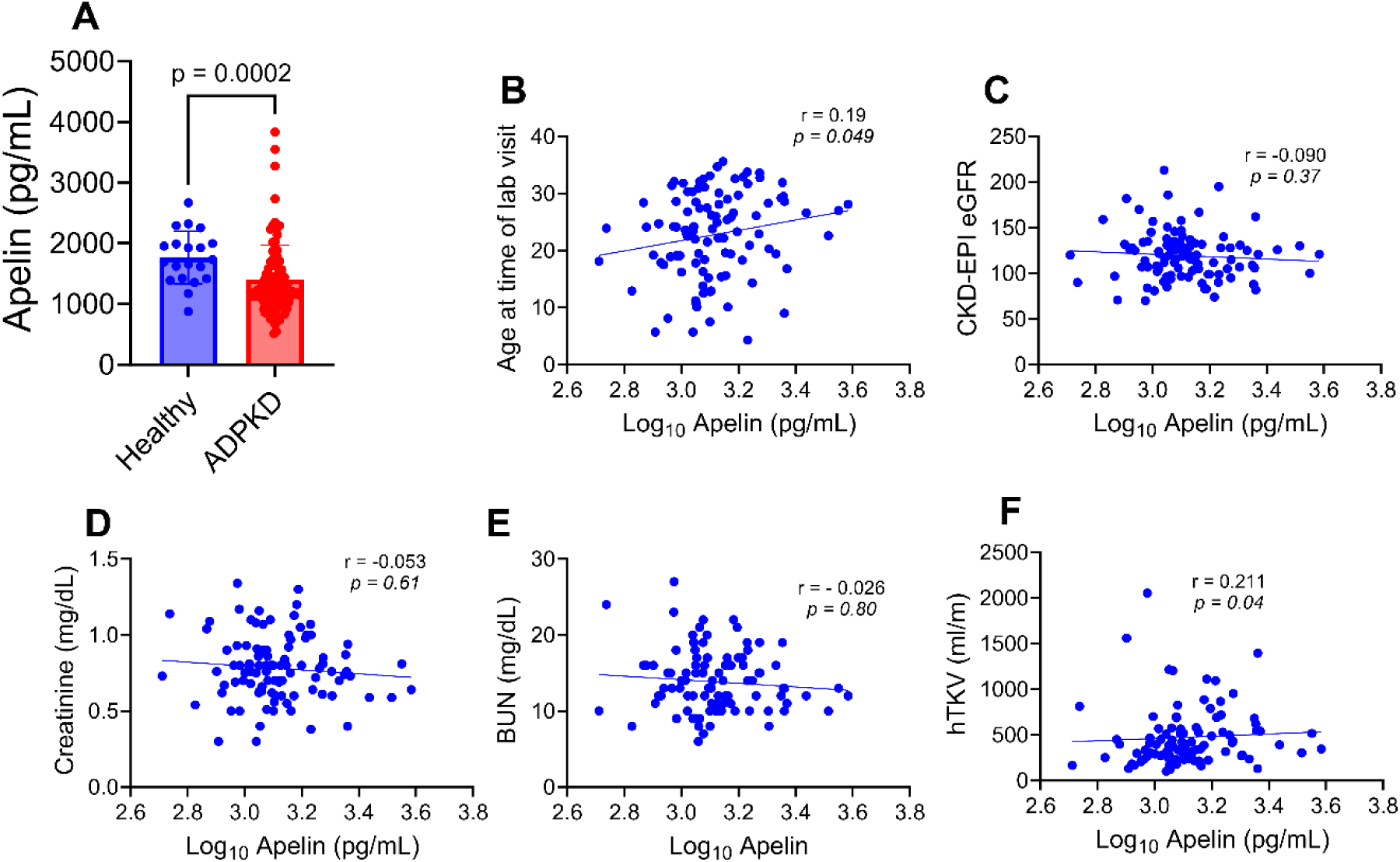
Circulating levels of apelin peptides in age- and sex-matched human ADPKD patients and healthy controls. (A) plasma apelin level, (B) correlation analysis of age of subjects at lab visit and plasma apelin, (C) correlation analysis of CKD-EPI eGFR and plasma apelin, (D) correlation analysis of creatinine and plasma apelin, (E) correlation analysis of blood urea nitrogen (BUN) and plasma apelin and (F) correlation analysis of heighted-adjusted total kidney volume (TKV) and plasma apelin. Healthy normal controls, n = 20; ADPKD patients, n = 100. Bar graphs represent mean±SD.

**Table 1.** Baseline clinical characteristics of ADPKD patients and normal controls in the EPOC study.

|  |  | Participants |  |
| --- | --- | --- | --- |
|  |  | Healthy Subject<br>(n=20) | ADPKD (n=100) |
| Sex, M/F |  | 7/13 | 39/61 |
| Age |  | 26.1±5.5 | 22.8±7.5 |
| BUN (mg/dL) |  | 12.65±3.6 | 13.8±4.1 |
| Creatinine (mg/dL) |  | 0.87±0.17 | 0.79±0.22 |
| eGFR (mL/min/1.73m<sup>2</sup>) |  | 104.8±17.1 | 119.4±27.0 |
| hTKV (mL/m) |  | - | 479±336 |
| MIC |  |  |  |
|  | A | - | 12 |
|  | B | - | 19 |
|  | C | - | 26 |
|  | D | - | 23 |
|  | E | - | 20 |
| PKD gene |  |  |  |
|  | PKD1 | - | 87 |
|  | PKD2 | - | 6 |
|  | Uncertain | - | 7 |
| Mutation type |  |  |  |
|  | Non-Truncating | - | 26 |
|  | Truncating | - | 65 |
|  | Uncertain | - | 9 |

MIC = Mayo imaging classification

### Effect of apelin on in vitro cyst growth of human ADPKD cells

Various concentrations of [Pyr^1^]apelin-13, a peptide ligand of the apelin receptor, were tested on the proliferation of human ADPKD cells using MTT assay. We found that 1 µM [Pyr^1^]apelin-13 decreased ADPKD cell proliferation by ∼20%, compared to control media (Fig. S3). [Pyr^1^]apelin-13 and azelaprag, a small molecule apelin receptor agonist, were also tested on *in vitro* cyst growth of human ADPKD cells cultured within 3D collagen matrix. Forskolin, a cAMP agonist, and epidermal growth factor (EGF) were added to induce cyst growth (Fig. 2A-E). Both [Pyr^1^]apelin-13 and azelaprag caused a dose-dependent reduction in cyst volume in the presence of forskolin and EGF. Azelaprag, but not [Pyr^1^]apelin-13, reduced the number of cysts per well (Fig. 2C-E).

**Figure 2.**
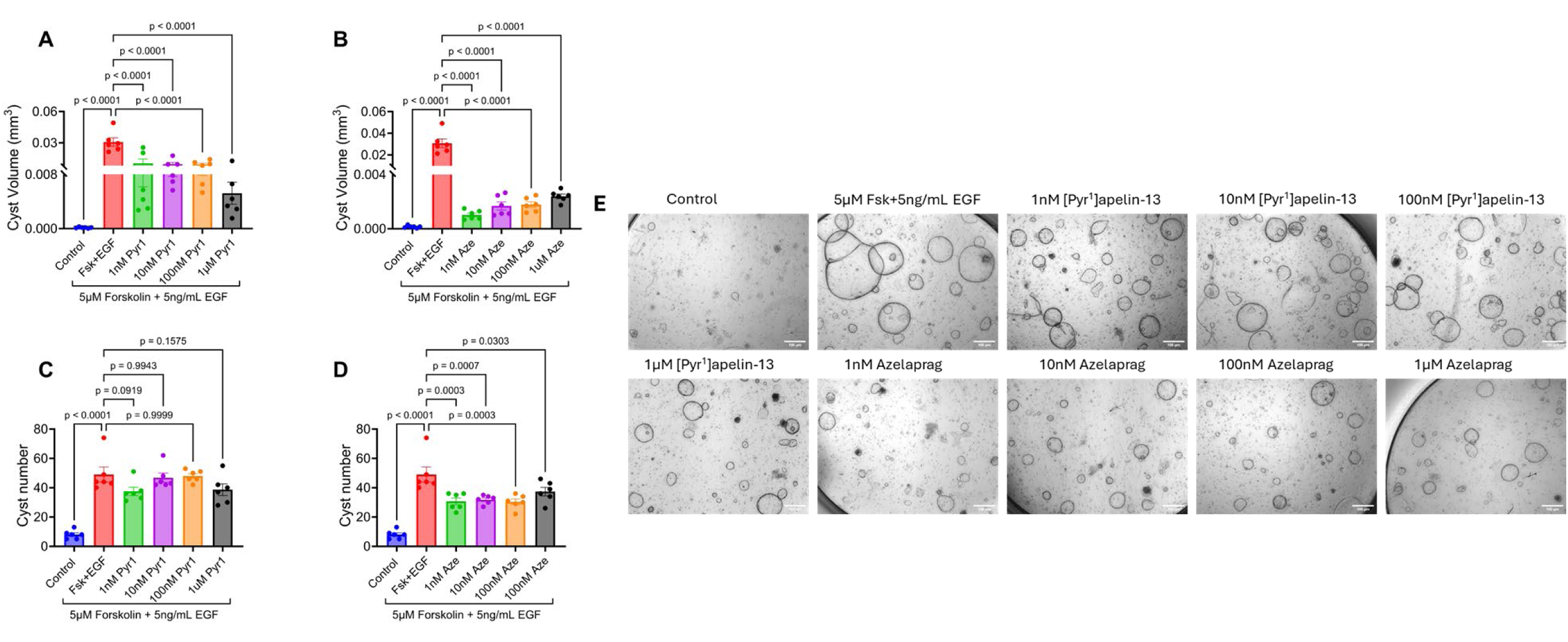
Apelin receptor agonists, [Pyr^1^]apelin-13 and azelaprag inhibited cyst growth in 3D model of primary human ADPKD cells. (A) effect of [Pyr^1^]apelin-13 on cyst volume, (B) effect of azelaprag on cyst volume, (C) effect of [Pyr^1^]apelin-13 on cyst number, (D) effect of azelaprag on cyst number, (E) images of 3D cysts treated with apelin or azelaprag. N = 6 per group. Bar graphs represent mean±SEM.

### Effect of apelin receptor agonists on PKD progression in mice

To test the hypothesis that exogenous apelin receptor agonism would inhibit cyst growth in PKD, we randomized *Pkd1^RC/RC^:Pkd2^+/-^* (PKD) mice, an orthologous early-onset PKD model, to receive vehicle, [Pyr^1^]apelin-13, azelaprag, or mozavaptan (a V_2_R antagonist used as a positive control) (Fig. S4). Mice were treated from postnatal day 22-48, a time period when cysts were rapidly growing and has been successfully used to show therapeutic inhibition of cyst growth^30^. Afterwards, the mice were killed and kidneys and heart were collected for evaluation. We found that daily treatment of PKD mice with [Pyr^1^]apelin-13 significantly reduced the two kidney weight-to-body weight ratio (KW/BW), similar to that observed with mozavaptan (Fig. 3A-B). [Pyr^1^]apelin-13 and mozavaptan also decreased cystic index and cyst number (Fig. 3C-D). By contrast, azelaprag did not affect %KW/BW or cystic index, but it significantly reduced cyst number (Fig. 3C-D). BUN was significantly reduced by treatment with mozavaptan and [Pyr^1^]apelin-13, while azelaprag had no effect (Fig. 3E). There were not changes in heart weight-to-body weight ratio in the PKD mice compared to control mice, and effect due to the treatment with the apelin receptor agonist or mozavaptan (Fig. 3F).

**Figure 3.**
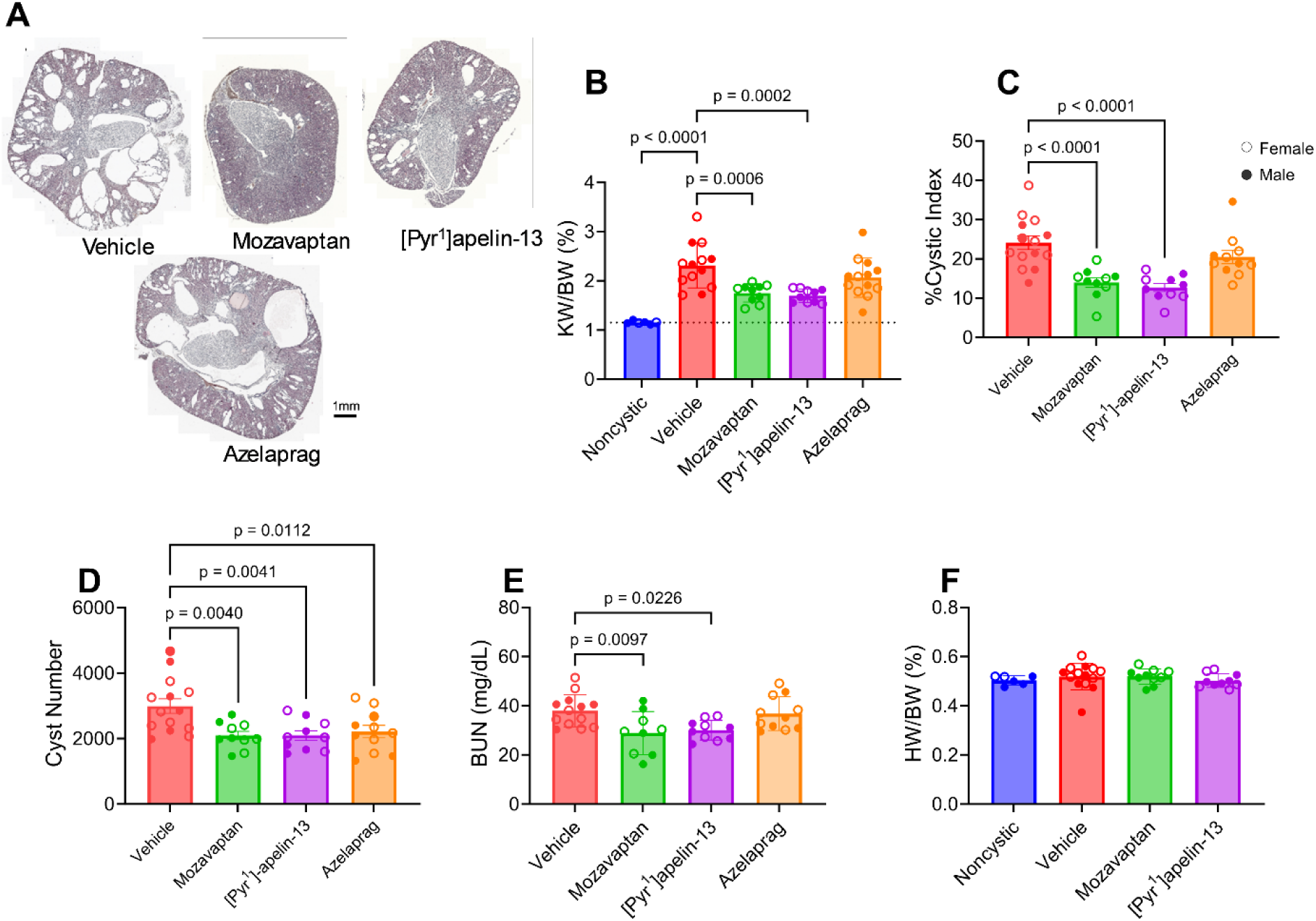
Effects of exogenous apelin receptor agonist, [Pyr^1^]apelin-13 or azelaprag on cyst growth in mice. (A) representative histological images of kidneys, (B) percent two kidney weight to body weight ratio (%2KW/BW), (C) percent cystic index, (D) cyst number, (E) blood urea nitrogen (BUN), (F) percent heart weight to body weight (%HW/BW). Phenotypic normal (noncystic), n=6; vehicle or azelaprag, n=13/group, mozavaptan or [Pyr^1^]apelin-13, n=10/group. Open cycles represent female animals. Bar graphs represent mean±SEM.

### Effect of apelin receptor agonists on renal cAMP and ERK activation

Activation of apelin receptors stimulate inhibitory G proteins, thereby inhibiting cyclic AMP production. Previously, we found that cAMP stimulates ERK-dependent cell proliferation, an important feature of renal cyst growth^4,38^. We assayed cAMP and phosphorylated ERK (pERK1/2) levels in kidney homogenates from PKD mice treated with [Pyr^1^]apelin-13, mozavaptan, azelaprag, or vehicle, and phenotypic normal (noncystic) mice. Renal cAMP levels were significantly increased in PKD mice compared to noncystic control animals, and this elevation in cAMP was significantly reduced by treatment with mozavaptan and [Pyr^1^]apelin-13, but not by azelaprag (Fig. 4A). Immunoblot analysis revealed that treatment with [Pyr^1^]apelin-13 and mozavaptan significantly decreased the level of pERK1/2 in PKD kidneys, demonstrating the reduction in ERK activity (Fig. 4B).

**Figure 4.**
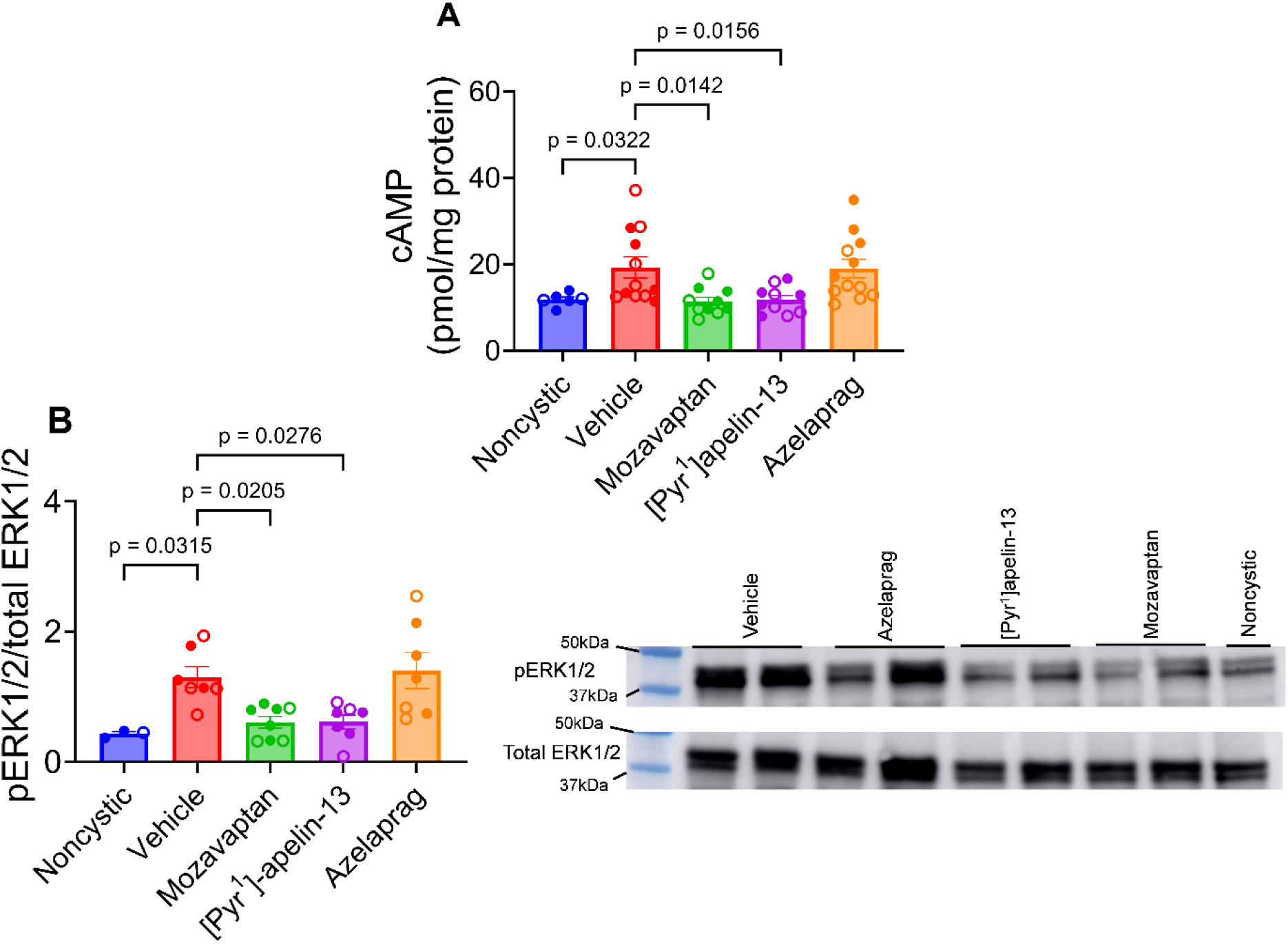
[Pyr^1^]apelin-13 inhibited cAMP production and ERK1/2 signaling in cystic kidneys. (A) kidney cAMP concentration, (B) phosphorylated ERK1/2/total (pERK1/2) relative to total ERK1/2 and a representative western blot image (right panel). Noncystic, n=6; Vehicle (n=7 for WB) or azelaprag, n=13 each, mozavaptan or [Pyr^1^]apelin-13, n=10 each. For western blot: noncystic (n = 3); vehicle, (n = 7), azelaprag (n = 7), mozavaptan (n = 8), [Pyr^1^]apelin-13 (n = 7). Bar graphs represent mean±SEM.

### Effects of apelin receptor activation on inflammation and the renin-angiotensin pathway in cystic mice

In PKD mice, inflammation was significantly increased as measured by the mRNA levels of key inflammatory markers, *Ccl2, Tnfα,* and *Il1β* expression, compared to noncystic control mice. Treatment with mozavaptan or azelaprag did not significantly affect the levels of these markers (Fig. 5A-C). However, [Pyr^1^]apelin-13 treatment significantly reduced *Il1β* mRNA expression and showed a trend towards reduced expression of the other inflammatory markers. We also investigated the effect of [Pyr^1^]apelin-13 treatment on fibrosis in these kidneys. However, neither [Pyr^1^]apelin-13 nor Mozavaptan affected renal fibrosis in this model, possibly because there was no significant fibrosis at this age of the PKD mice (Fig. S5A-B).

**Figure 5.**
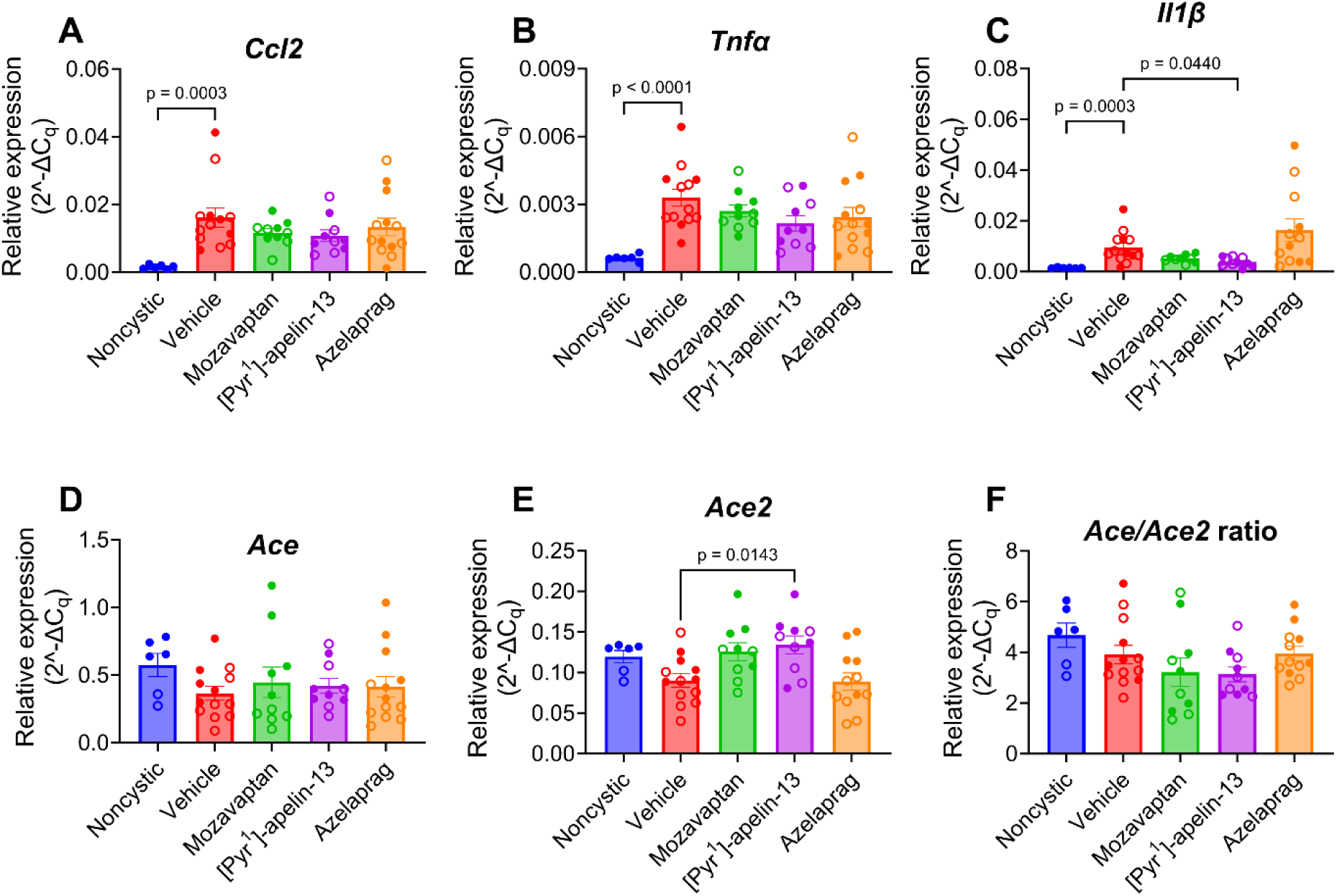
Effect of exogenous [Pyr^1^]apelin-13 treatment on inflammation and RAS pathway in the kidneys. A, *Ccl2*, B, *Tnfα*, C, *Il1β*, D, *Ace*, E, *Ace2,* F. *Ace/Ace2* ratio. Noncystic, n = 6; Vehicle (n = 7 for WB) or Azelaprag, n = 13 each, Mozavaptan or [Pyr^1^]apelin-13, n = 10 each.

In heart disease, one mechanism of cardioprotection mediated by the apelin signaling pathway involves inhibition of the renin-angiotensin system (RAS)^20,39^. We wondered whether the disease-modifying effects of the apelin system in PKD involved modulation of the RAS and measured the mRNA levels of *Ace* and *Ace2*, as well as their ratio, in the kidneys. Treatment with mozavaptan or azelaprag did not affect the mRNA levels of *Ace* or *Ace2* (Fig. 5D-F). [Pyr^1^]apelin-13 caused a slight, but significant, increase in the mRNA of *Ace2* without affecting *Ace* or the *Ace/Ace2* ratio (Fig.5F).

### Effect of apelin on metabolic and extracellular matrix markers in PKD mice

To further understand the disease-modifying effects of apelin signaling in PKD, we performed bulk RNA sequencing on vehicle- and apelin-treated kidneys and set the adjusted *p-value* at 0.05 and the fold change at >2. Based on these criteria, we identified 20 significantly upregulated genes, including *Acaa1b, Hcar1, Dio1, and Cy26b1* (Fig. 6A). We found 195 downregulated genes, including *Lcn2 (Ngal), Mmp14, Col1a1, A2m, Postn, Havcr1 (Kim-1), and Dcdc2*, all of which had previously been linked to increased PKD progression^40–42^. We selected two upregulated (*Dio1* and *Acaab1)* and two downregulated (*Timp1* and *Mmp14*) genes for confirmation by qPCR using Taqman assays (Table S1). In selected samples for RNAseq analysis and the whole population, *Timp1* was significantly downregulated (Fig. 7A, Fig. S6A). Although *Mmp14* was downregulated, this was not significant (Fig. 7B, Fig. S6B;). Both *Dio1* and *Acaab1* were significantly upregulated in the selected samples used for RNAseq and across the entire dataset (Fig.7C-D, Fig. S6C-D).

**Figure 6.**
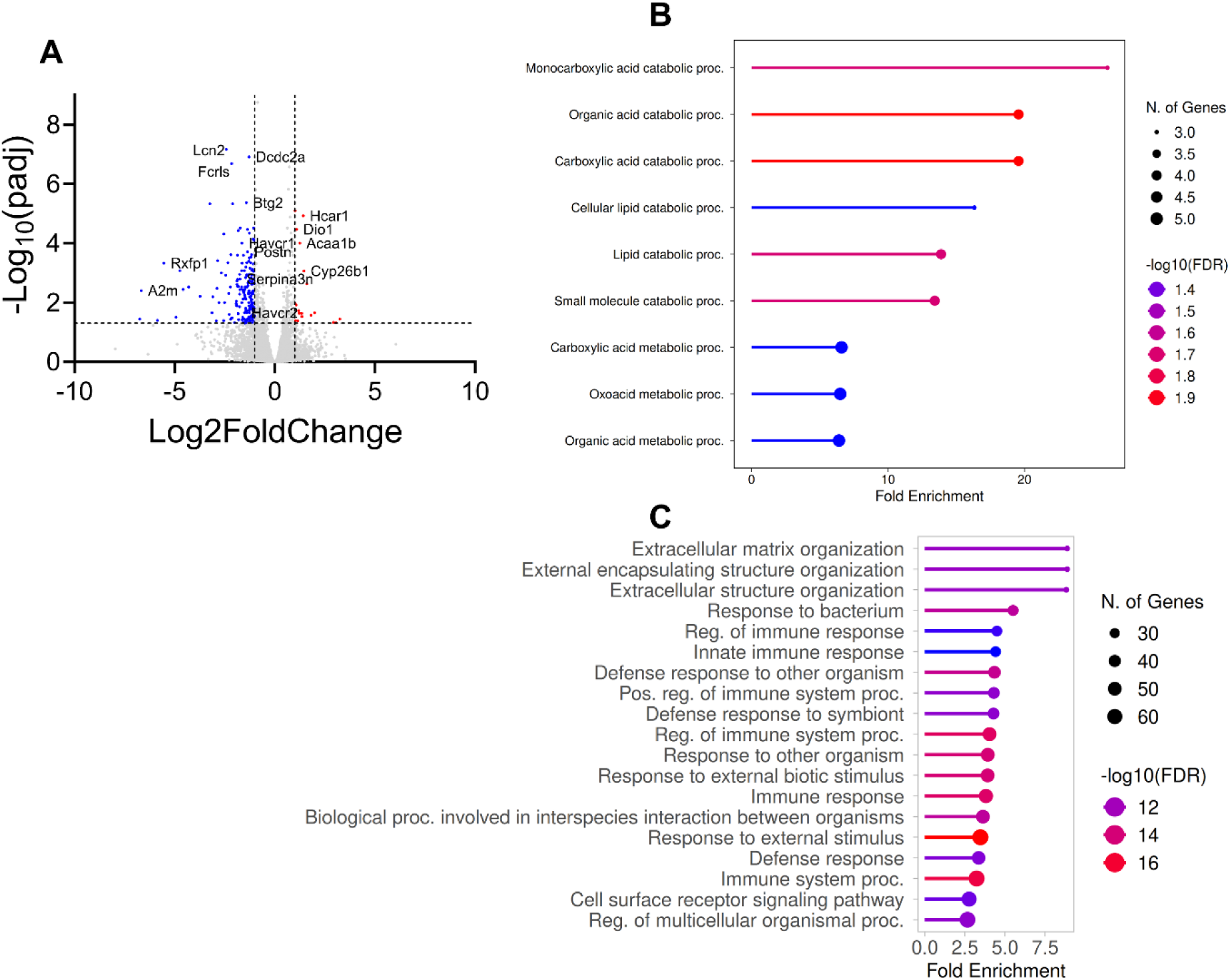
Differentially expressed genes between vehicle- and apelin-treated PKD mice. (A) volcano plot of differentially expressed genes in apelin-treated relative to vehicle-treated mice, (B) dot plot showing top enriched biological process pathways in gene ontology that were upregulated by Pyr^1^]apelin-13 treatment compared to vehicle, (C) dot plot showing top enriched biological process pathways in gene ontology that were downregulated by Pyr^1^]apelin-13 treatment compared to vehicle. N = 3 samples per group.

**Figure 7.**
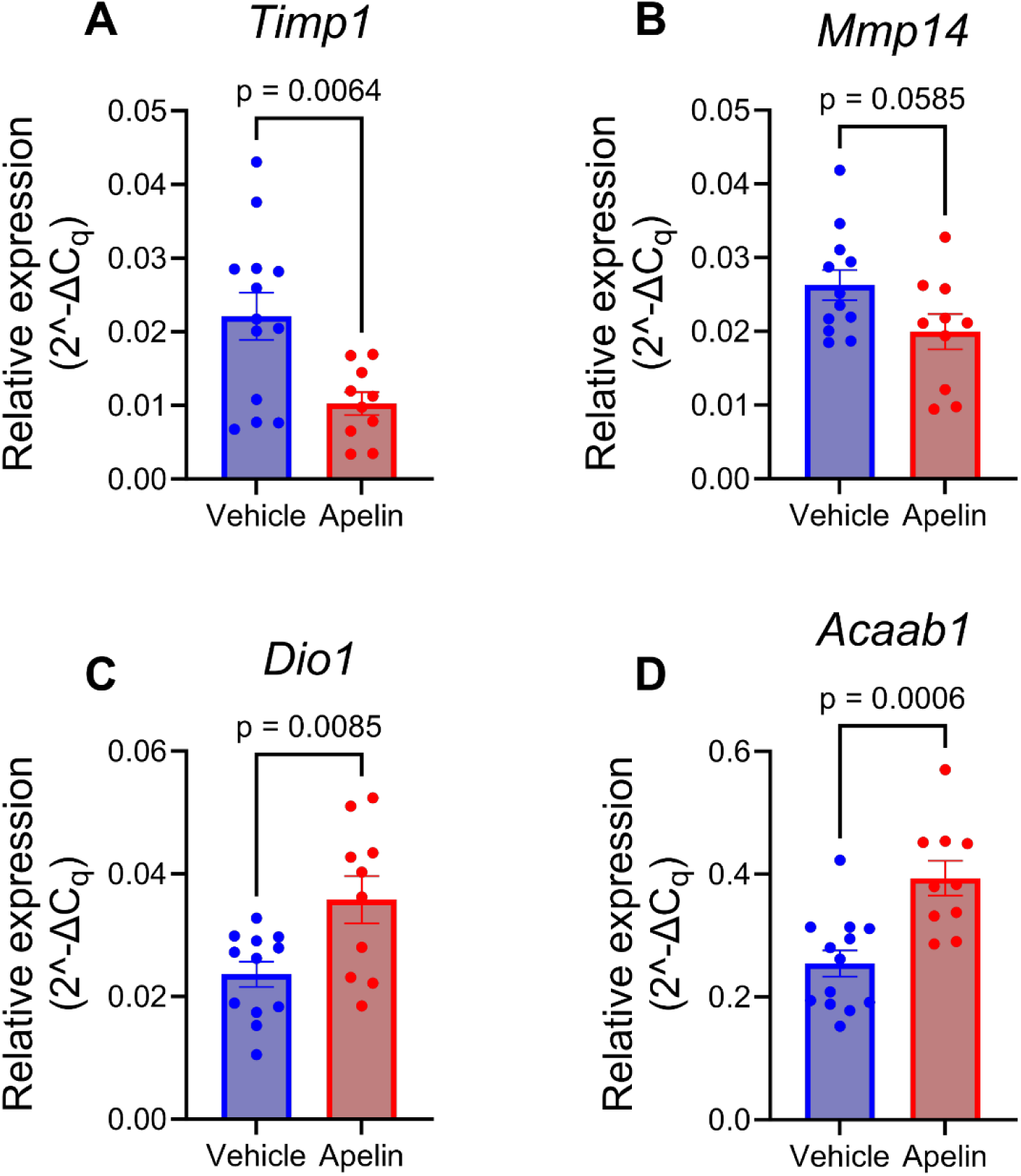
pPCR validation of differentially downregulated (A-B) or upregulated (C-D) genes identified from bulk RNAseq data. (A) *Timp1*, (B) *Mmp14*, (C) *Dio1,* and (D) *Acaab1*. Noncystic, n = 6; Vehicle or azelaprag, n = 13 each, mozavaptan or [Pyr^1^]apelin-13, n = 10 each.

Biological process pathway analysis of these genes revealed that the top upregulated pathways were associated with carboxylic acid/organic acid metabolism and cellular lipid metabolism (Fig. 6B). Whereas the top downregulated pathway genes were involved in extracellular matrix structure and organization (Fig. 6C). Furthermore, we compared our data with those previously published by Song *et al.*^43^ and Swenson-Fields *et al.*^44^ to evaluate how our set of differentially expressed genes aligned with their findings using the Ingenuity Pathway Analysis (QIAGEN IPA) software. Interestingly, in both datasets, the alterations in upstream regulators of PKD were highly consistent with those observed in our own analysis (Fig. S7).

### Effect of apelin receptor agonists on urinary output in mice

One of the major side effects of V_2_R inhibition with tolvaptan is excessive diuresis due to impaired concentrating capabilities of the kidneys. Hence, tested whether apelin receptor activation has the same adverse effect since both signaling pathways modulate intracellular cAMP production. In wild-type animals, we did not observe any effect of [Pyr^1^]apelin-13 on urine volume (diuresis) or osmolality at 2 mg/kg and 4 mg/kg body weight (Fig. 8A, D). At the lower dose (2 mg/kg BW), azelaprag did not affect urine volume or osmolality, whereas the higher dose significantly increased urine volume without affecting osmolality (Fig. 8B, E). Mozavaptan significantly increased in urine volume (diuresis) (Fig. 8C) and reduced urine osmolality (Fig. 8F)

**Figure 8.**
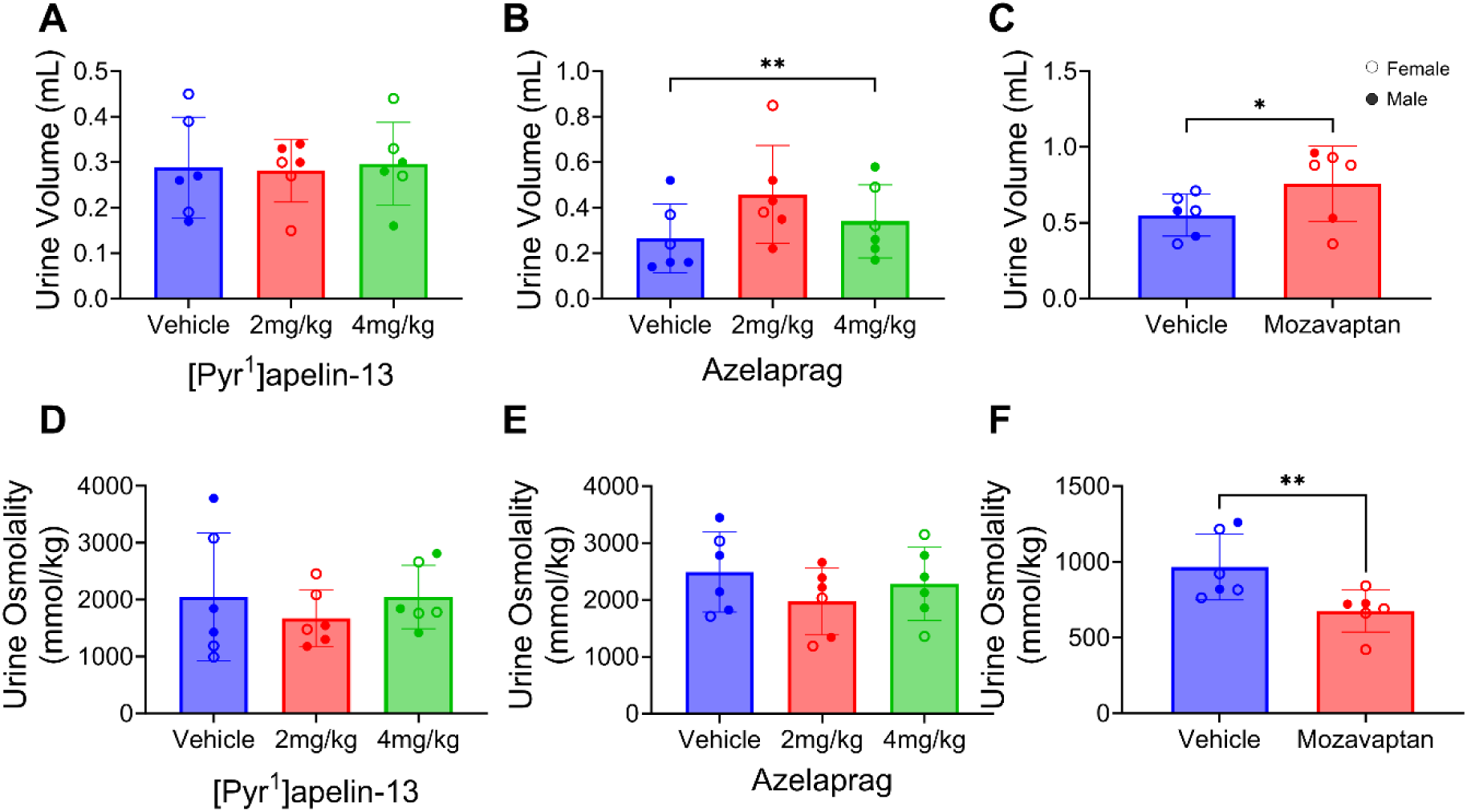
[Pyr^1^]apelin-13 did not promote diuresis or urine osmolality in wildtype mice. Urine volume for: (A) [Pyr^1^]apelin-13, (B) azelaprag, (C) mozavaptan; Urine osmolality for: (D) [Pyr^1^]apelin-13, (E) azelaprag, (F) mozavaptan. 0.9% saline was used as vehicle for [Pyr^1^]apelin-13 and azelaprag while vehicle for mozavaptan was water. N = 6. Bar graphs represent mean±SEM.

## Discussion

In this study, we report that circulating apelin levels were significantly reduced early in ADPKD patients, despite markers of renal function (BUN, creatinine, and eGFR) being completely normal, suggesting that circulating apelin could be a novel prognostic biomarker. We also showed that activation of the apelin receptor with apelin inhibited cyst growth in 3D microcyst cultures of primary human ADPKD cells and in an orthologous mouse model of PKD, and improved kidney function (measured by BUN). These affects appeared to be mediated by inhibiting cAMP production and reducing ERK1/2 signaling, a key signaling pathway in cyst growth^4^. RNA-seq analysis demonstrated that apelin’s disease-modifying effects in PKD involved downregulation of genes previously shown to promote cyst growth, including *Timp1, Lcn2*, and *Mmp14,* and upregulation of genes involved in lipid metabolism, including *Acaab1*^40–42^. Importantly, activation of the apelin receptor did not promote diuresis or alter urine osmolality in mice, suggesting that the mechanisms mediating apelin’s effects in PKD differ from those of V_2_R inhibition.

The apelin signaling pathway primarily signals via Gαi-mediated inhibition of cAMP production^12,21,45^. Based on this, we would expect activation of apelin receptor signaling to mirror the diuretic effect of a V_2_R antagonism. However, our data suggest that inhibition of cyst growth, at the apelin dose used in the study (Fig. 8), was mediated via a different mechanism, or combination with cAMP inhibition. Importantly, disruptions of intracellular Ca^2+^ signaling and/or reductions in intracellular Ca^2+^ levels have long been thought to contribute to the aberrant cell proliferative phenotype characteristic of ADPKD cells^4^. Perhaps apelin receptor agonism modulates both Gαi and Gαq signaling in ADPKD cells resulting in cAMP inhibition and intracellular calcium release, as previously described in adipocytes^46^. Additionally, although apelin receptor action was suggested to promote diuresis^12,47,48^, these studies were conducted in lactating animals, marked by a significant increase in the synthesis and secretion of AVP to preserve water for maximal milk production^47^, and our experiments were conducted in wildtype male and female (non-pregnant) animals. Hence, apelin treatment may avoid the excessive urination and thirst associated with taking tolvaptan in ADPKD patients, possibly by activating the Gαq signaling pathway together with or different from cAMP or via Gαi proteins that work on specialized pools of cAMP not affected by AVP, such those in primary cilia^49,50^, given the mechanosensitive functions of the apelin receptor^51^.

Surprisingly, azelaprag inhibited cyst growth *in vitro*, but did not have the corresponding effect in the PKD mice, at least at the tested dose. The higher concentration (4 mg/mL) of this small molecule led to a increase in urine volume without altering urine osmolality, despite not having any effect on urine osmolality or diuresis at the concentration tested (2 mg/mL) in the PKD mouse model. This may suggest that both azelaprag and apelin engage distinct downstream signaling pathways upon receptor binding. For instance, apelin binding to the apelin receptor is known not only to stimulate Gαi-mediated inhibition of cAMP production but also to activate Gαq signaling^15,46,52^, whereas azelaprag is known to engage only Gαi signaling^21^. Alternatively, the differences in response to the endogenous peptide apelin and the small molecule agonist, azelaprag, could be in their biodistribution. The endogenous peptide apelin readily distributes to the kidneys and collects in the bladder^53^, whereas azelaprag first passes through the liver where it is metabolized and released back into circulation^24^. The concentration of azelaprag that ultimately reaches the kidney is unknown but could be too low to elicit a therapeutic response in the PKD mice. This may also explain why doubling its dose in the diuretic experiment produced some effect on urine volume (Fig. 8B), as well as the lower dose did not affect urine output. Incidentally, human trials of azelaprag as an obesity drug were recently terminated due to liver damage^54^. Taken together, our data are consistent with the hypothesis that the disease-modifying responses to apelin involve the engagement of multiple downstream signaling pathways involved in cystogenesis, inflammation and fibrosis in PKD mice, as evidenced by the RNA-seq data.

The observation that apelin receptor activation inhibited cyst growth in 3D microcyst assays using primary human ADPKD cell cultures, as well as *in vivo* in an orthologous mouse model of PKD, is very promising. Mechanistically, we showed that the disease-modifying role of apelin in PKD was mediated by inhibition of cAMP synthesis and ERK1/2 activation, although our RNA-seq data suggest other, as-yet unidentified pathways, are also likely involved. This is important, as there remains an unmet clinical need for novel therapeutic strategies that are safe and effective while avoiding the negative consequences of tolvaptan. Interestingly, our data suggest that apelin receptor activation is devoid of the negative effects observed with tolvaptan.

Additionally, the major cause of death in ADPKD patients following the introduction of renal replacement therapy is cardiovascular disease, including hypertension. Hence, given the vasodilatory (blood-pressure-lowering) and anti-cardiac hypertrophic effects of the apelin system^7,10,36^, targeting the apelin receptor is likely to represent a promising therapeutic avenue for ADPKD patients. Moreover, several clinical studies have examined the potential of targeting the apelin receptor in other diseases, including pulmonary hypertension^55^ and, more recently, chronic kidney disease^56^, and have found this target to be safe and well-tolerated. Thus, our data may provide the requisite support to test apelin agonists in ADPKD patients.

One limitation of testing apelin peptides in humans is its extremely low plasma stability, as well as the requirement for administration by injection. Unfortunately, the only small-molecule apelin receptor agonist to have advanced furthest in clinical development is azelaprag. Hence, further studies will be required to screen small-molecule compound libraries for novel apelin receptor agonists that could be developed for the treatment of ADPKD. Additionally, one weakness of this study is that we did not test different doses of the agonists to identify the optimal concentration for maximal response. Another weakness is that we only tested apelin in a juvenile mouse model of PKD that does not progress to renal failure; therefore, additional studies will be required to evaluate the long-term effect of apelin on PKD progression.

In conclusion, we demonstrated that circulating levels of apelin were significantly reduced in children and young adults with ADPKD before detectable changes in kidney function and that exogenous apelin reduced cell proliferation and cyst growth of primary human ADPKD cells and renal cystic disease in an orthologous PKD mouse. We propose that the apelin receptor represents a potential novel therapeutic target for the treatment of ADPKD.

## Acknowledgement

This work was funded by a grant from the PKD Foundation (D. N) and NIH/NIDDK (R01DK115727 to A.Y.). Human ADPKD cells were provided by the PKD Biospecimens and Biomaterials Core of the Kansas PKD Center (U54 DK126126 to A.Y., S.P., and D.W.), and the PKD Research Resource Consortium. We thank Sarah Tague from the KUMC imaging core (supported by the KIDDRC (NIH U54 HD 090216)) for her help with imaging and quantification of the cysts. We also thank the University of Kansas Medical Center - Genomics Core for their help with RNA extraction, library preparation, and bulk RNA sequencing. The Genomics Core is supported by NIH S10 High-End Instrumentation Grant (NIH S10OD036343) and Frontiers CTSA grant (UL1TR002366).

