## Supplementary file for "Apelin inhibits cyst growth and improves kidney function in mice with polycystic kidney disease"

**Detailed method**

Human Studies

Plasma samples collected were collected in EDTA tube from age- and sex-matched healthy subjects and ADPKD patients through the multicenter Early PKD Observation Cohort (NCT02936791). Briefly, patients included in this cohort were between ages 4 and 35 years old, diagnosed with ADPKD through imaging or genetic testing, and had an estimated glomerular filtration rate (eGFR) >80mL/min per 1.73m^2^. All healthy subjects included in this study had no family history of kidney disease, normal eGFR, and came from all races/ethnic groups. The exclusion criteria included non-insulin or insulin-dependent diabetes mellitus, inability to provide written informed consent, systemic illness (including lupus erythematosus, vasculitis), and unavailability for magnetic resonance imaging (MRI) and blood/urine collections.

**Histology, cystic index, and blood urea nitrogen measurements**

Kidneys were cut transversely into two halves, processed, and sectioned at the Histology Core at the University of Kansas Medical Center. Sections (5 µm thick) were deparaffinized, rehydrated, and Masson Trichrome Staining (Cat. No. NC0017745; Polysciences, Warrington, PA ) performed according to the manufacturer's protocol. Kidney sections were automatically imaged on a Nikon Eclipse High Content Analysis (HCA) system (Tokyo, Japan), comprised of a Ti-E inverted motorized microscope. The HCA/JOBS function of Nikon NIS AR software was used to automatically load the slides, find the tissue, autofocus, and acquire large, tiled images of each kidney section with a Nikon Plan Apo 2x (NA 0.2) objective using an 8-bit Nikon DS-Fi3 color camera. General Analysis was used to automatically threshold each image using the red channel to create binary masks that covered the entire kidney section (Red 2-210) or cysts (Red 195-255, size >10 µm), these thresholds were then combined to ensure that all cyst thresholded areas in an image limited to the area occupied by the total kidney section threshold, as previously described^1^. The binaries for each image were inspected and, if needed, edited with the binary editor. This included removing the outer papillary space from the cyst threshold. Automated measurement results from the edited whole kidney and cyst binaries were processed in Excel to determine cystic index per section (two sections per mouse). For comparison, cystic index was also measured manually for a subset of the sections as previously described using ImageJ.

**Cell culture and proliferation Assay**

The use of de-identified surgically discarded tissues complies with federal regulations and is approved by the Institutional Review Board at KUMC. Primary human ADPKD cells were obtained from the Kansas PKD Research and Translational Core Center, and the Polycystic Kidney Disease Research Resource Consortium (PKD RRC). Cells were cultured in DMEM/F-12 supplemented with 5% FBS, 1% ITS (Cat No.: CB-40352, Corning™), and 1% penicillin-streptomycin (PenStrep) (Cat.# P0781, Sigma). Cells were used between passages 1 and 3 in all experiments.

For cell proliferation assay, 5x10^3^ cells were seeded per well of a 96 well plate in complete media and changed to 0.002% FBS media 24hrs later. After another 24hrs of serum-starvation, cells were treated with different concentrations the apelin for 48hrs. Epidermal growth factor (EGF), 25ng/mL, was used as positive control. Cell proliferation was then performed using Cell counting reagent (CCK8) kit according to manufacturer’s protocol and data normalized to vehicle (PBS)-treated controls.

**RNA sequencing and data processing**

Total RNA was extracted from tissue samples obtained from six mice, comprising three apelin-treated and three vehicle-treated animals. mRNA libraries were prepared from total RNA using the Tecan Universal Plus mRNA Kit, which employs poly(A) selection for transcript enrichment. Libraries were sequenced on an Illumina NovaSeq X Plus platform to generate paired-end reads (101 bp × 2). Each biological replicate was sequenced across two lanes, producing paired-end FASTQ files for each lane. All downstream analyses were carried out using R (v4.5.0) within RStudio (v2026.01.0), with statistical analyses performed using Bioconductor packages and external command-line tools employed for quality control and transcript quantification.

Raw sequencing reads were assessed for quality using FastQC (v0.12.1), and quality metrics were aggregated across samples using MultiQC (v1.33). Read quality assessment indicated high base quality scores and no evidence of excess adapter contamination or systematic technical artifacts; therefore, no read trimming or filtering was applied. For each biological replicate, paired-end reads from the two sequencing lanes were concatenated to generate a single forward (R1) and reverse (R2) FASTQ file, ensuring that each sample was represented as a single sequencing library for downstream analyses.

**Transcript quantification and gene-level count estimation**

Transcript-level abundance estimation was performed using Salmon (v1.10.1) in quasi-mapping mode. A transcriptome index was constructed from the protein-coding transcripts of the GENCODE mouse reference annotation (release vM38). Salmon was applied to each merged paired-end FASTQ file to estimate transcript abundances, generating transcript-level quantification files. Transcript-level abundance estimates were imported into R using the tximport package (1.38.2). Transcript abundances were summarized to the gene level using a transcript-to-gene mapping file derived from the same GENCODE reference annotation. Gene-level counts were generated using length-scaled TPM values (countsFromAbundance = "lengthScaledTPM"), enabling appropriate normalization for transcript length while retaining compatibility with count-based differential expression methods.

**Identification of Differentially expressed genes and enrichment analysis**

Differential gene expression analysis was conducted using the DESeq2 package (v1.50.2). Gene expression was modeled as a function of treatment group (~ Group), with Vehicle-treated samples specified as the reference condition. Library size normalization and dispersion estimation were performed using DESeq2’s internal methods. Statistical significance was assessed using Wald tests, and p-values were adjusted for multiple hypothesis testing using the Benjamini–Hochberg false discovery rate procedure. Genes with an adjusted p-value < 0.05 were considered differentially expressed. To visualize the results, a volcano plot was generated using the was generated using ggplot2 (v4.0.1), with thresholds set at Log_2_foldchange (Log_2_FC)>1 and adjusted p-value (padj) < 0.05. Significant genes were highlighted by mapping differential expression status to point color, and the y-axis was transformed to -log_10_(padj) for statistical visualization. Functional enrichment of DEGs was performed via Gene Ontology (GO) pathway analysis using cluster Profiler (v5.2.0), utilizing the org.Mm.eg.db (v3.21.0) mouse genome database.

**Table**

**Table S1. Taqman gene expression probes and assay ID used for RT-PCR experiments.**

| Gene | Assay ID |
| --- | --- |
| *Ezr* (housekeeping gene) | Mm00447761_m1 |
| *Ace2* | Mm01159006_m1 |
| *Ace* | Mm00802048_m1 |
| *Il1β* | Mm00434228_m1 |
| *Tnfα* | Mm00434228_m1 |
| *Ccl2* | Mm00434228_m1 |
| *Acaab1* | Mm00728805_s1 |
| *Dio1* | Mm00839358_m1 |
| *Mmp14* | Mm00485054_m1 |
| *Timp1* | Mm01341361_m1 |

**Results**

**Figures**

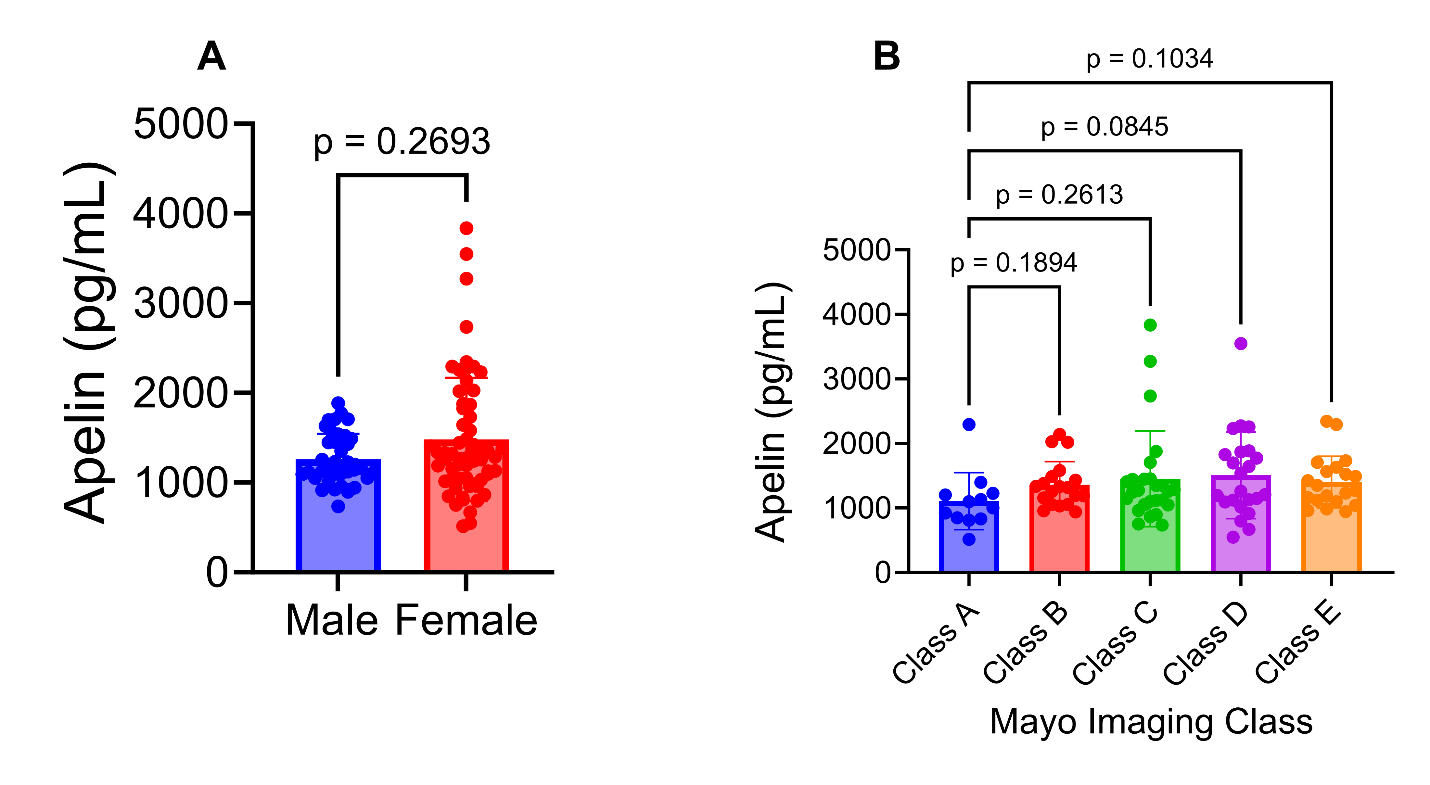

Figure S1. Comparison of circulating apelin levels in ADPKD patients. A, circulating levels of apelin in male and female ADPKD patients; Male, n = 39; female, n = 61.; B, circulating levels of apelin grouped based on Mayo Imaging Classification. Class A (n=12), Class B (n=19), Class C (n=26), Class D (n=23), and Class E (n = 20). Data represents mean±SD.

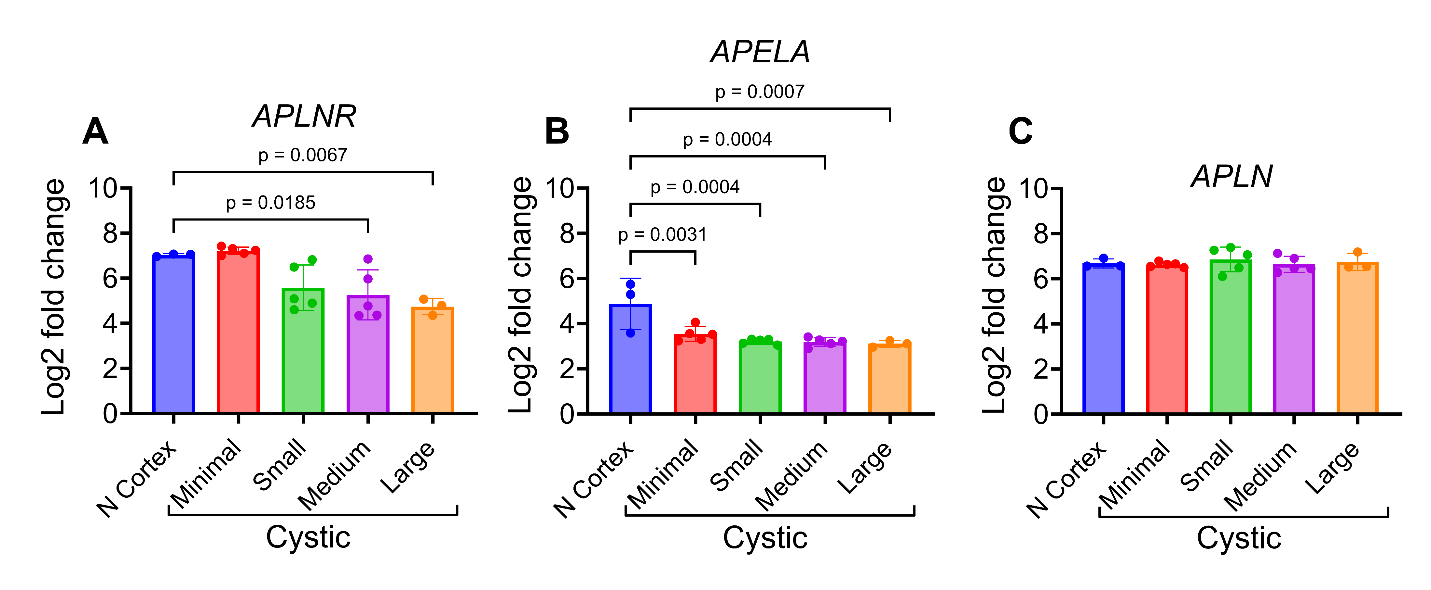

Figure S2. Apelin signaling pathway is dysregulated in human ADPKD. Reanalysis of the public dataset, GSE7869 from global gene expression profiling study of renal cysts reported by Song et al.^2^. The microarray expression data was obtained from renal cysts of various sizes from 5 ADPKD patients (*PKD1* mutations) and 3 kidneys (cortex) from healthy subjects. Data represents mean±SD.

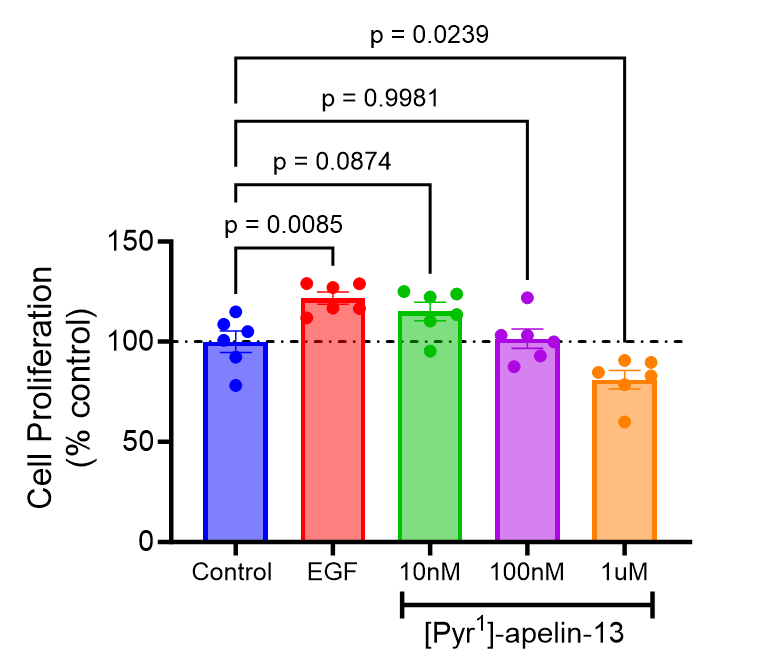

Figure S3. Apelin inhibited proliferation of primary human ADPKD cells. Epidermal growth factor (EGF) was used at 25ng/mL used as positive control. Data represents mean±SEM.

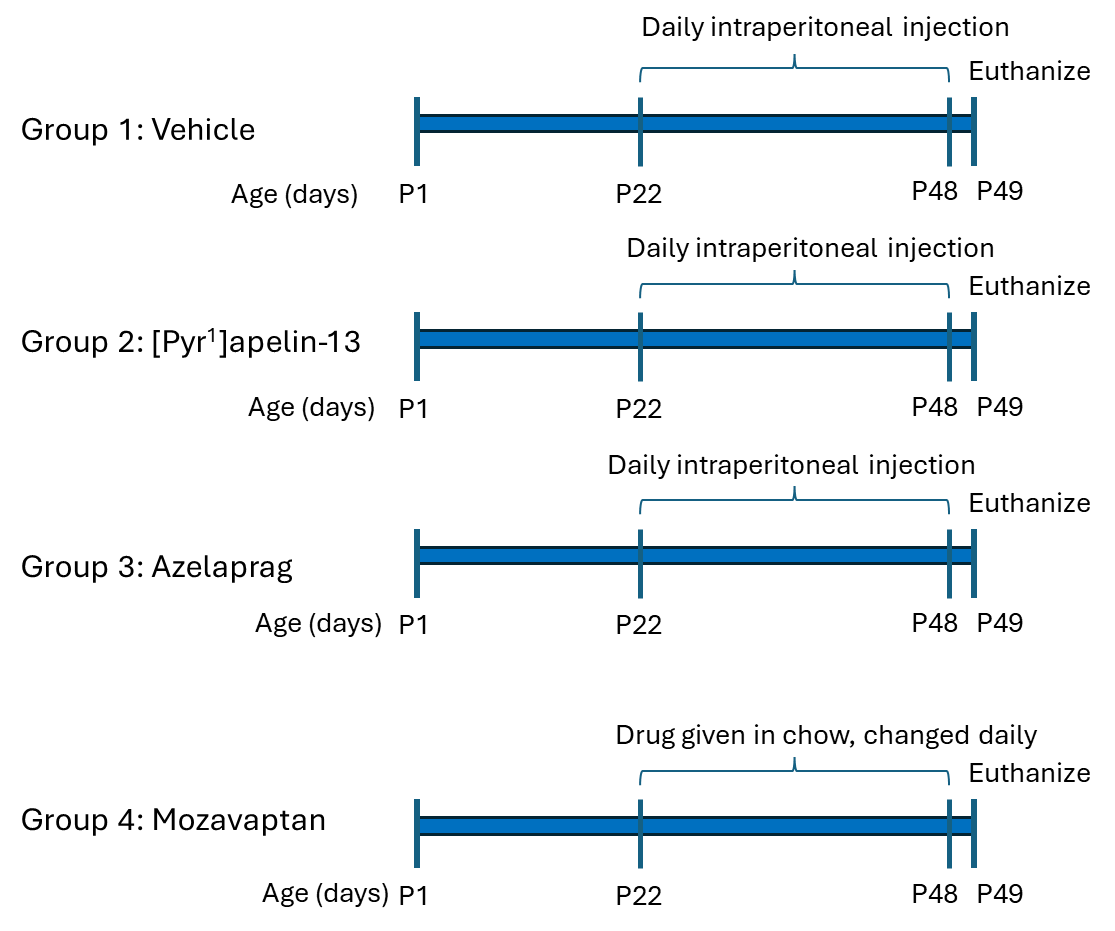

Figure S4. *In vivo* treatment schedule for *Pkd1^RC/RC^:Pkd2^+/-^* mouse model. Both [Pyr^1^]apelin-13 and azelaprag (apelin receptor agonists) were given at 2 mg/mL, while mozavaptan was given in powdered chow and replaced daily.

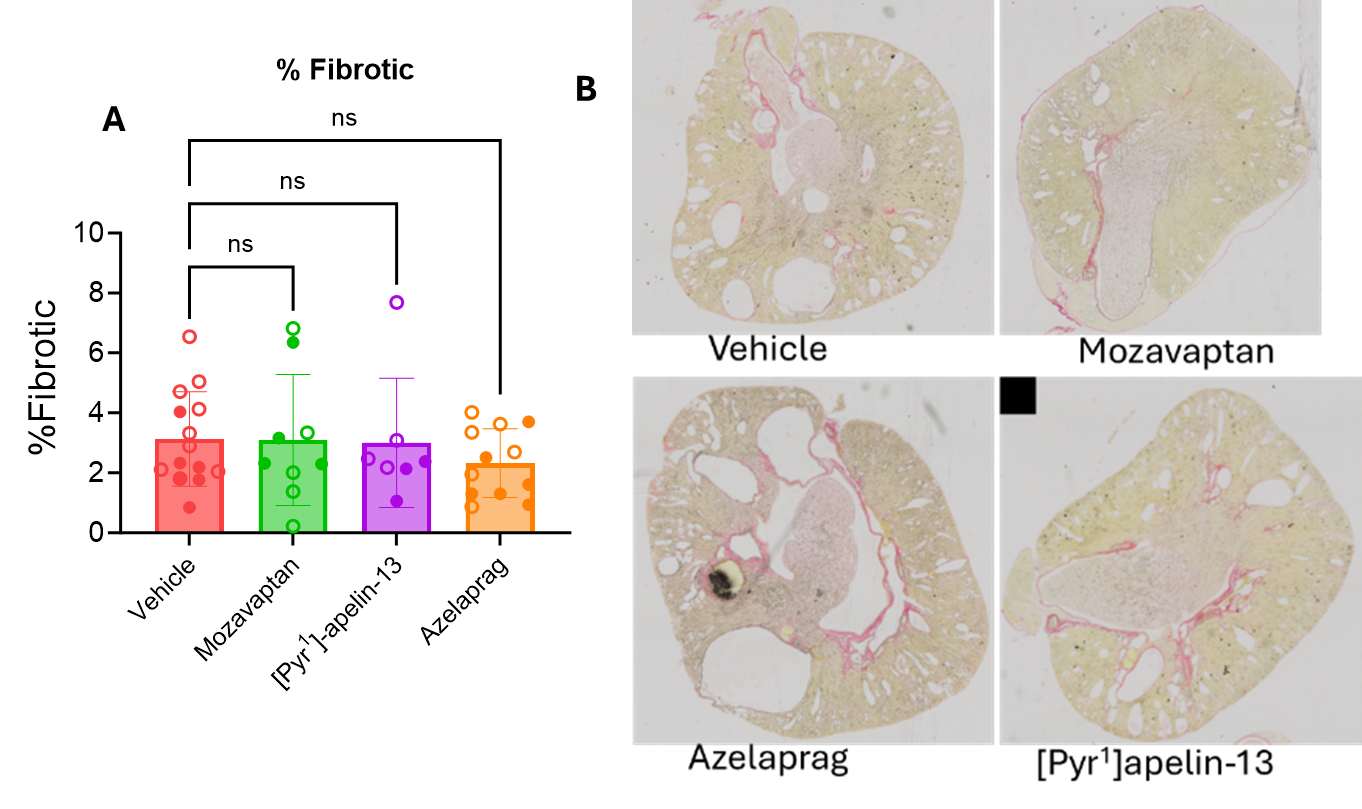

Figure S5. Effect of exogenous [Pyr^1^]apelin-13 treatment on fibrosis in cystic mice. A, quantification of picro Sirius red staining of collagen in the PKD mouse kidneys, B, representation micrograph of PKD mouse kidneys. % fibrosis was determined from the total area staining over the total kidney area.

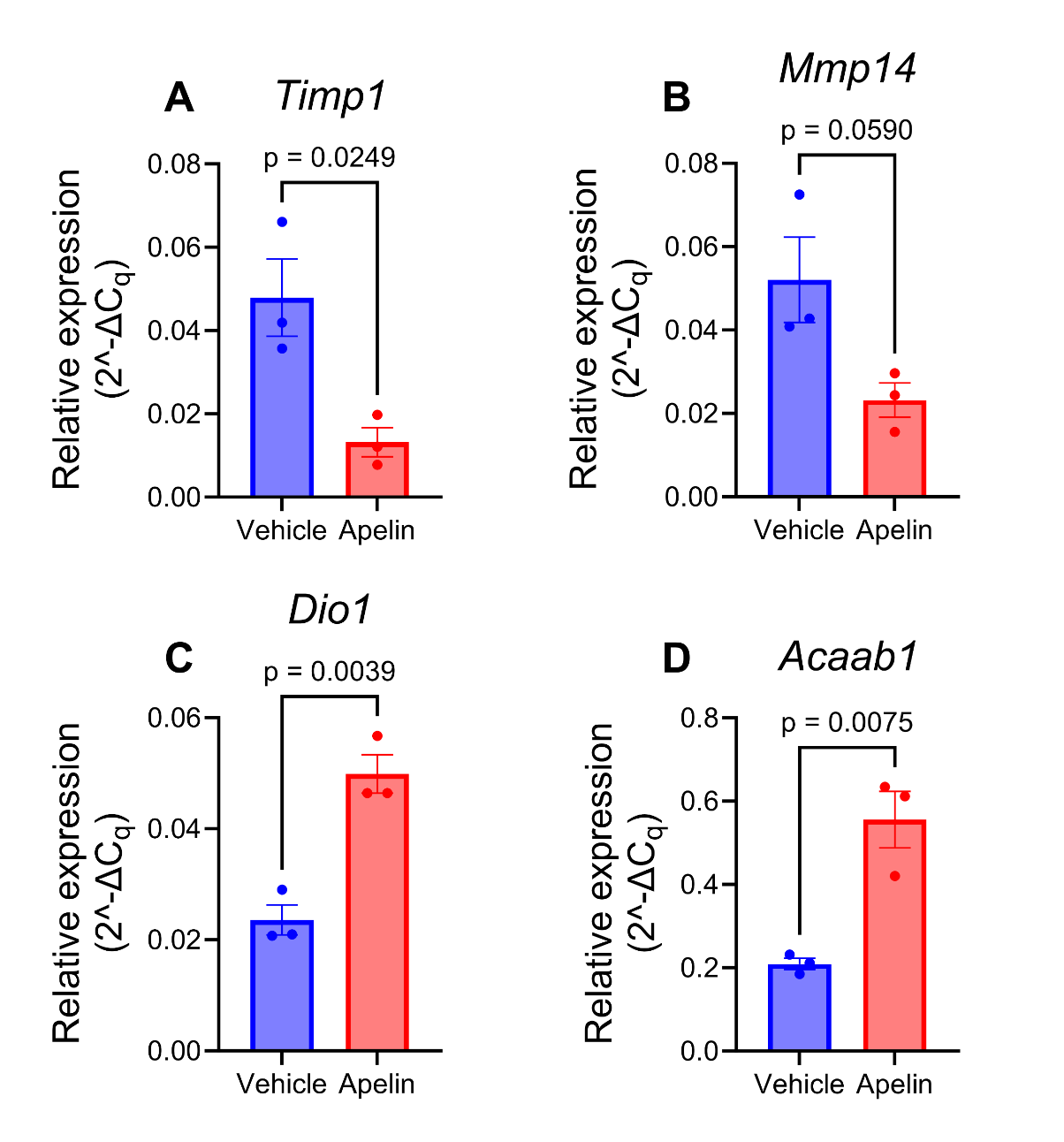

Figure S6. pPCR validation of differentially downregulated (A-B) or upregulated (C-D) genes identified from bulk RNAseq data. A, *Timp1*, B. *Mmp14*, C. *Dio1* and D. *Acaab1*. Vehicle, n = 3, [Pyr^1^]apelin-13, n = 3.

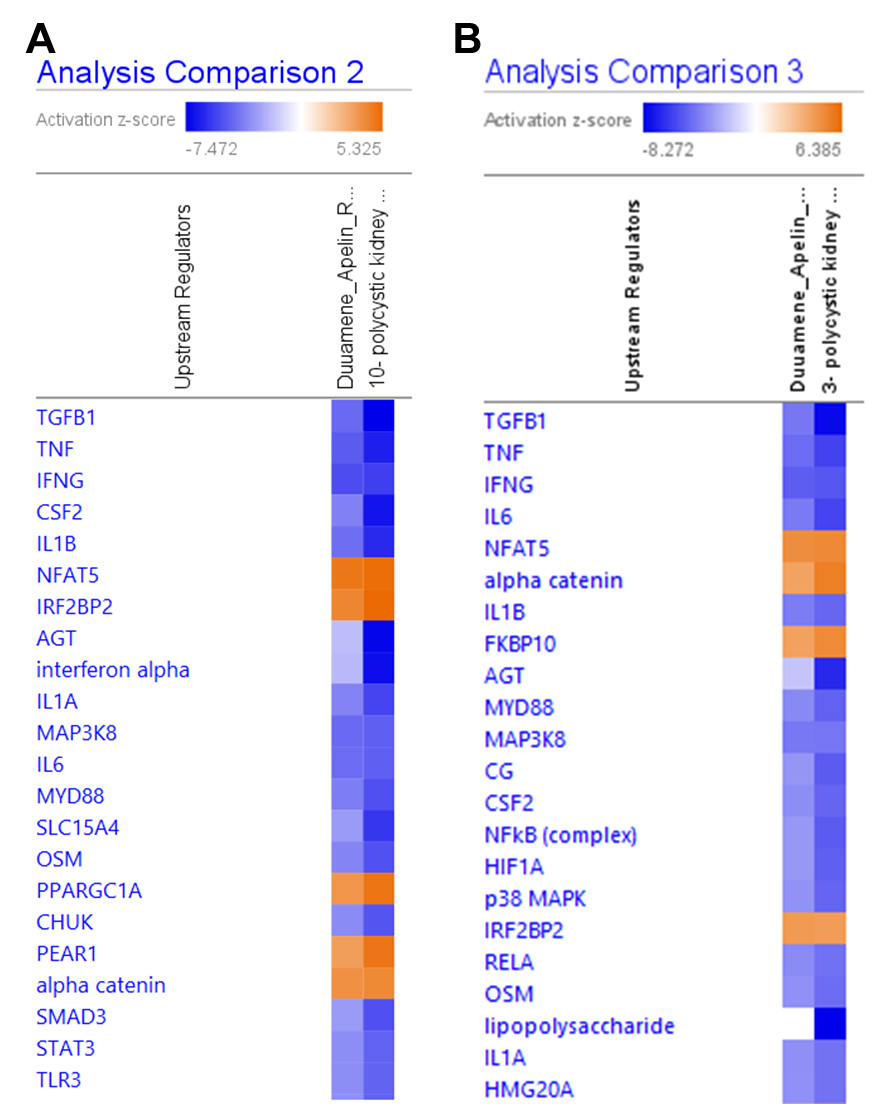

Figure S7. Upstream regulators of PKD common between our dataset and published datasets. A., Dataset published by Song et al.^3^; B., Dataset published by Swenson-Fields et al.^4^ The QIAGEN IPA heatmap uses activation z-scores to predict the functional state of upstream regulators, with orange indicating predicted activation and blue indicating predicted inhibition. These scores were calculated by comparing our experimental gene expression changes against the QIAGEN Knowledge Base, a massive library of established biological relationships. Thus, if our data aligns with the literature's expected direction of change for a regulator's targets, a significant z-score (>2 or <-2) is generated. The software decides which regulators to display based on a p-value of overlap (0.05).
